# The frequent occurrence and metabolic versatility of *Marinifilaceae* bacteria involved in organic matter mineralization as a key member in global deep sea

**DOI:** 10.1101/2022.07.25.501493

**Authors:** Jianyang Li, Chunming Dong, Qiliang Lai, Guangyi Wang, Zongze Shao

## Abstract

Transfer of animal and plant detritus of both terrestrial and marine origins to the deep sea occurs on the global scale. Microorganisms play an important role in mineralizing them therein, yet to identify *in situ*. Here we report the family *Marinifilaceae* that occurred as one of the most predominant bacteria thriving on the new inputs of plant and animal biomasses in both marginal and oceanic areas observed via *in situ* incubation about their differentiation, environmental adaption, and metabolic mechanisms underlying their prevalence in organic matter mineralizing communities. We described the metabolic features and *in situ* metabolizing activities of different subgroups (tentative genus level), based on the metagenomic and metatranscriptomic data. One representative subgroup MF-2 dominated plant detritus-enriched cultures and specialized in polysaccharide degradation by encoding many hydrolases involved in the hydrolysis of hemicellulose, pectin, starch, cellulose, and polysaccharides containing N-acetyl groups; this subgroup also encodes a manganese superoxide dismutase with the potential of lignin oxidation and possesses complete nitrogen fixation pathway to compensate for the shortage of nitrogen sources inside the plant detritus. In contrast, those dominating the animal tissue-supported microbiomes were more diverse and formed three subgroups, which distinguished themselves from MF-2 in carbon and nitrogen metabolisms. Regardless of differentiation in carbon and nitrogen metabolisms, they share in common in energy conservation through organic fermentation, and anaerobic respiration of diverse electron receptors. These results highlight the role of *Marinifilaceae* bacteria neglected before in organic matter mineralizing in marine environments coupling carbon and nitrogen cycling with metals and other elements.

**IMPORTANCE:** Microbial mineralization of organic matters has a significant impact on the global biogeochemical cycle. This report confirmed the role of *Marinifilaceae* in organic degradation in the oceans, with underestimated contribution in the ocean carbon cycling. It is the dominant taxon thriving on plant and animal biomasses in our *in situ* incubator, as well as in whale- and wood-falls. At least nine subgroups were revealed, and widely distributed in global oceans but merely predominant in organic-rich environments with an average relative abundance of 8.3 %. Different subgroups display a preference for the degradation of different macromolecules (polysaccharides, lignin, and protein) and adapt themselves to the environments via special metabolic metabolisms.

## INTRODUCTION

Deep sea, the largest dark biota on earth (1, 2), plays a major role in the global material and elements biogeochemical cycles (3). The principal source of carbon and energy for the dark sea biota is the primary production from the upper layer of the ocean and the resulting deposition of particulate organic matter (POM), dissolved organic matter (DOM), and refractory dissolved organic carbon (rDOM) via multiple pumps (4, 5). The organic matters (OMs) input into deep sea, including POM, DOM, and rDOM, critically contribute to the establishment of habitats for deep-sea microbial communities (6).

Heterotrophic microorganisms play a keystone role in the mineralization of those OMs in the deep sea, particularly the dissolution of POM to DOM and subsequent absorption of cleavage products, both of which are considered rate-limiting steps of deep-sea microbial metabolism (7). The particle-associated microorganisms in the deep sea have been examined by filter fractionation techniques and sediment traps (7–14). The sediment trap study indicated the central role of piezophile-like Gammaproteobacteria and Epsilonproteobacteria in the mineralization of sinking POM in the deep sea (13). Another report showed that particle-associated microorganisms, such as Sphingomonadales, Rhodobacterales, and Alteromonadales, play a primary role in the cleavage of organic matter by releasing their extracellular enzymes into the particles in the bathypelagic realm (14).

In comparison, little was known about the microbial-mediated mineralization of the large and fast sinking particles, such as oceanic or terrestrial animal and plant tissues like well-known ecosystems of wood-fall, kelp-fall, and whale-fall (15–19). The remains of animal and plant input into the deep sea occur on the global scale every year continuously, which are regarded as “hot times” and “hot spots”. Those large and fast sinking particles may escape disaggregation, dissolution, and solubilization en route to the deep sea (20), and impact the microbial communities in the deep sea (21, 22). Around 500 Tg C of terrestrial OM was imported into the ocean annually and ca.10% is buried in marine sediments (23, 24). Although its amount is less than POM and DOM both derived from primary production in the sunlit surface waters of the ocean, the impact of terrestrial OM input into the deep sea cannot be ignored (25), as well as the kelp-fall and whale-fall. Some studies have estimated that a single storm event could transport up to 1.8-4 Tg of driftwood carbon to the ocean and that a single large whale carcass could provide a flux equivalent to 2000 years of background POC flux, or 30-40 % of the annual carbon flux of a swarm of sinking swimming crabs to the deep ocean floor (26). Recently, a report has estimated that 4.7 million m^3^ of large wood could flow into the oceans each year (27). These estimates clearly demonstrate the importance of large organic falls to the ecology of deep-sea ecosystems. However, wood has been unheeded in the quantification of organic carbon burial on continental margins (28).

The degradation of OMs derived from those remains is a temporally dynamic process that contains the succession of specialized communities with distinct lifestyles and metabolic potentials in the deep sea (29). Localized, high organic concentrations may deplete oxygen, which in turn attracts anaerobic microbial communities thriving on the organic-falls (30, 31). These resulting habitats are described as chemosynthetic ecosystems because of the sulphide production finally inducing chemosynthetic communities (19, 32). However, their mineralization mediated by microbes remain less explored in deep sea, which hampers our ability to evaluate the fate and impacts of those bulky remains.

We hypothesized that the microorganisms thriving on those remains sunk into the deep-sea may be different from the bacteria observed in the deep water column sampled with CTD cassettes, filter fractionation techniques, and sediment traps, because large pieces rich in OM do not constantly sink through the water column and settle to the seabed at every location. To identify the bacterial assemblage involved in the *in situ* transformation and degradation of the fast and sinking POM in the deep sea, we performed deep-sea *in situ* incubation amended with various natural animal and plant tissues in different oceans (preprint: https://doi.org/10.21203/rs.3.rs-1700182/v1) (33). After 4-12 months of incubation via an *in situ* incubation device landing on the seafloor, we found that bacteria of the family *Marinifilaceae* (MF) were the predominant members of both animal and plant tissue enrichments. Whether and how this bacterial group plays a role in the depolymerization, degradation, and transformation of macromolecular organic matter remains uncertain. To date, only a few bacteria of this family have been isolated (34), such as *Labilibaculum manganireducens* (35), *Ancylomarina subtilis* (36), and *Marinifilum albidiflavum* (37), but distantly related to the ones in our *in situ* cultures. In this study, we analysed the genetic diversity, global distribution, environmental adaptation, and metabolic metabolism of MF via phylogenetic, metagenomic, and metatranscriptomic analyses. This knowledge will facilitate our understanding of its role in biogeochemical cycles in oceanic interiors.

## RESULTS AND DISCUSSION

### Diversity and wide distribution of MF bacteria in the global ocean

The neighbour-joining trees based on 16S rRNA gene sequences showed that the clones belonging to MF family were close to the genera of *Labilibaculum* and *Ancylomarina* (**Fig. 1A**) and that MF family contained at least 9 subgroups (tentative genus level), named MF-1 to MF-9 (**Fig. 1B**). Among the 9 subgroups, MF-2, MF-4, and MF-8 were separately closely related to the cultured bacteria of the genera *Labilibaculum*, *Ancylomarina,* and *Marinifilum*, while other subgroups have no pure cultures yet to match and might represent novel genera of this family. In particular, the MF bacteria detected in the five *in situ* enrichments were affiliated with the MF-1, MF-2, MF-3, and MF-4 subgroups.

**Figure 1.**
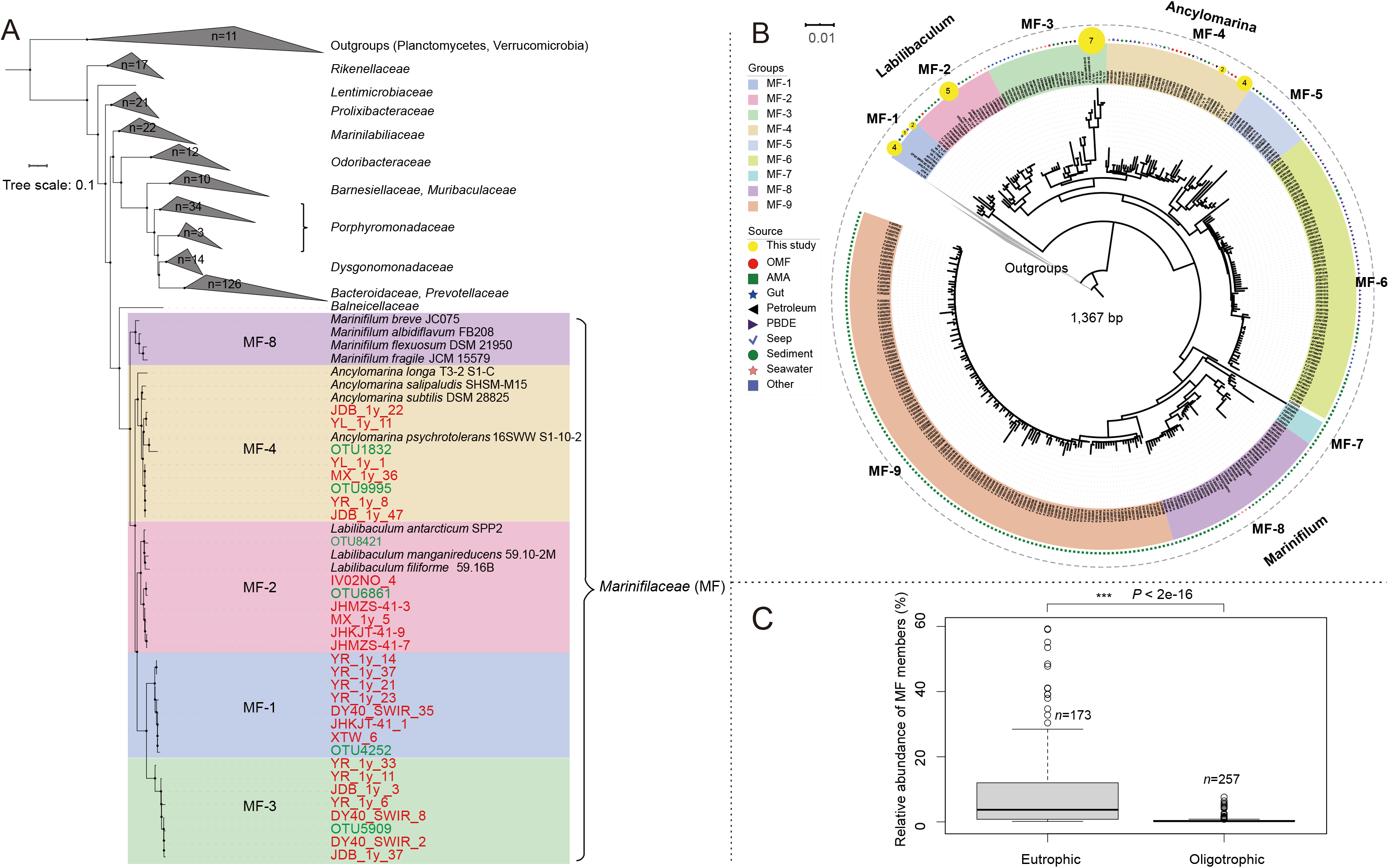
Phylogenetic relationships based on 16S rRNA sequences and comparative analysis of the relative abundance of MF members from different environments. (**A**) 16S rRNA gene phylogenetic tree containing 25 clones (marked red) and 6 OTUs (marked green) belonging to the MF family, as well as currently identified strains within Bacteroidales. Some strains within the phyla Planctomycetes and Verrucomicrobia were identified as outgroups. This tree shows the phylogenetic placement of MF members within Bacteroidales. (**B**) Phylogenetic tree based on the near-full-length 16S rRNA (1367 bp) gene sequences within the MF clade. There are at least nine subgroups within the MF clade. MF-2, MF-4, and MF-8 are separately closely related to the cultured bacteria of the genera *Labilibaculum*, *Ancylomarina*, and *Marinifilum*, while members of other subgroups have not been cultured individually yet, and these subgroups might represent novel genera of this family. The families *Prolixibacteraceae* and *Marinilabiliaceae* were the outgroups. OMF, organic falls; AMA, associated with marine animals; PBDE, polybrominated diphenyl ethers. (**C**) Variation analysis of the relative abundance of MF from the eutrophic and oligotrophic environments. Welch’s t test was used to estimate the significance among the samples, *P* ≤ 0.001 (***); the data used here are shown in Supplementary Table 1.

The global distribution pattern of MF-1, MF-2, MF-3, and MF-4 showed that they are widely distributed in marine environments, including seawater columns, marine sediments, marine animal and plant surfaces or animal intestines, and organic-rich marine habitats, such as whale-falls, wood-falls, seagrass detritus, cold seeps, and petroleum-contaminated sediment, but not found in terrestrial and freshwater environments (**Fig. S2**, **Table S1** and **Table S2**). Moreover, based on the metadata (**Table S1**), we found that the average abundance of the four subgroups in organic-rich environments, e.g., wood-falls, whale-falls, oil pollution, enrichments with seaweed or polysaccharide, was 8.3 %, which was significantly higher (*P* < 0.001) than that in oligotrophic sediment and seawater (**Fig. 1C**). Those results implied that MF bacteria are opportunistic bacteria that may exist as rare and ubiquitous species of famine waiting for the feast of sinking bulky POM in the deep sea, as suggested previously by Jorgensen (6).

### Preference of MF subgroups to different organic substrates

A total of 11 distinct metagenome-assembled genomes (MAGs) (bin B6 to bin B13, WF05, WF09, and WF13) belonging to MF family were obtained from the five metagenomic datasets and two wood-fall metagenomic datasets (19, 38) (**Table 1** and **Table S3**). They were respectively affiliated with the MF-1, MF-2, MF-3, and MF-4 subgroups in the phylogenomic tree (**Fig. 2B**), and may represent 10 different novel species based on the average nucleotide identity (ANI) values (**Table S4**).

**Table 1.**
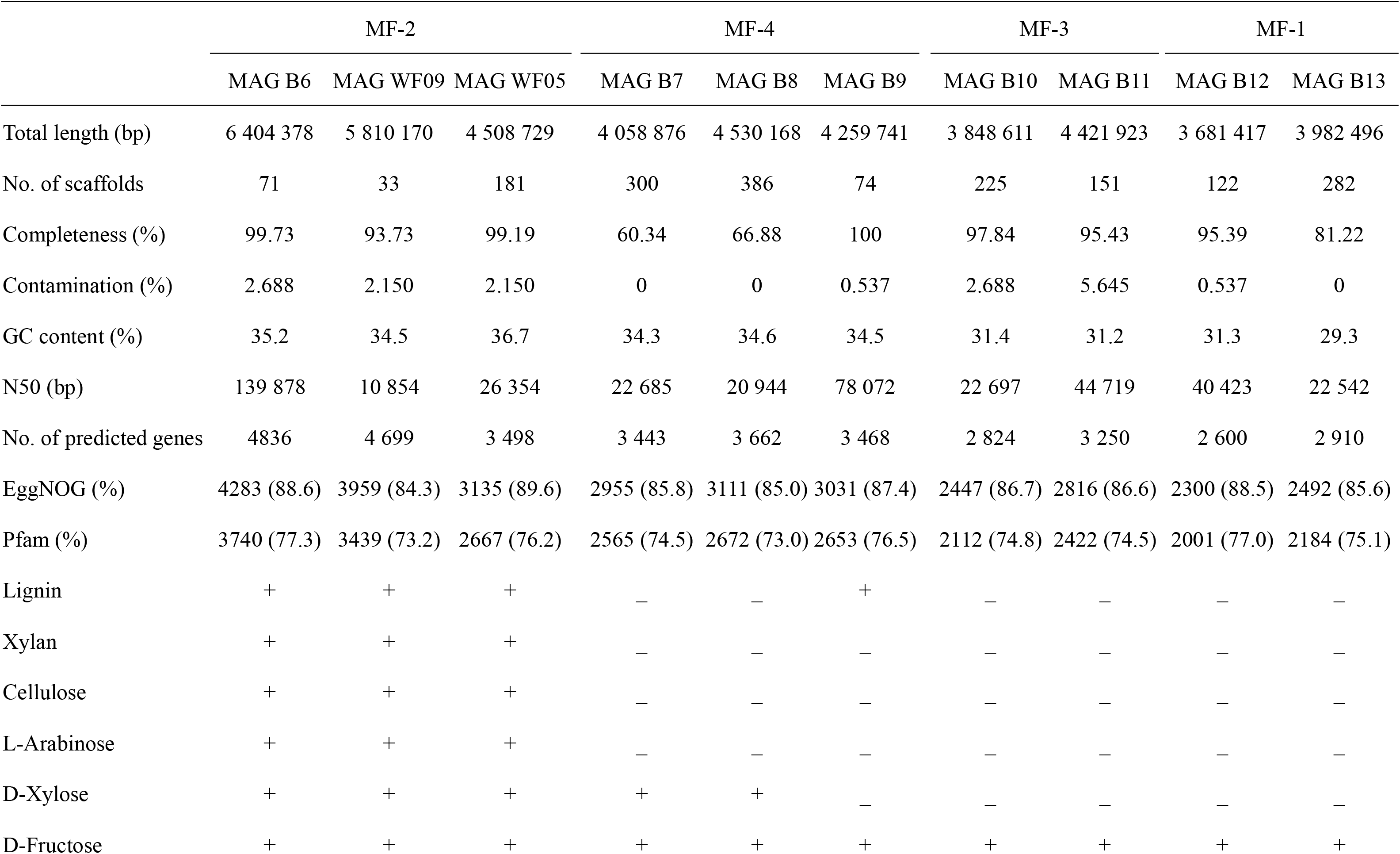
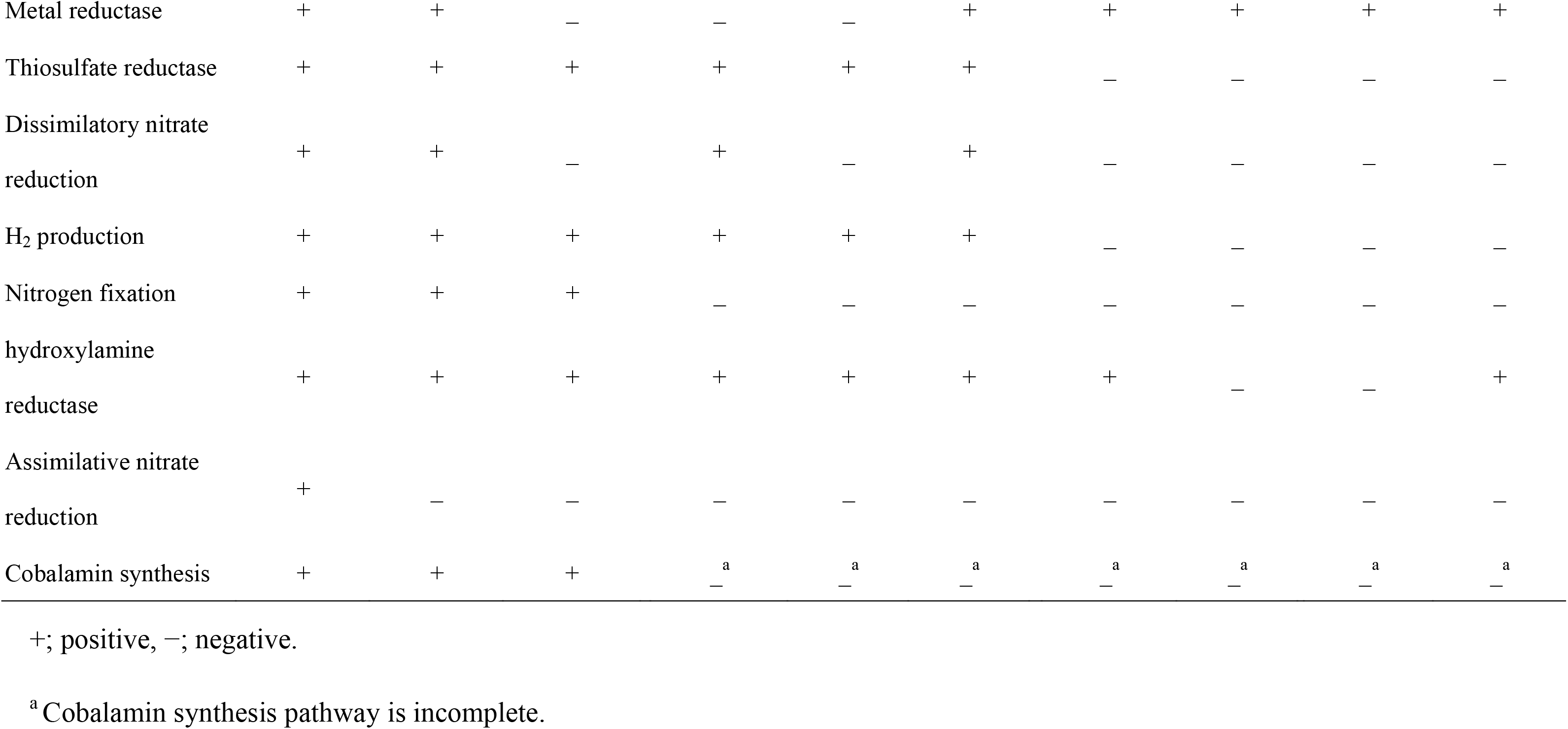
Genomic features and metabolic potential of MF MAGs.

**Figure 2.**
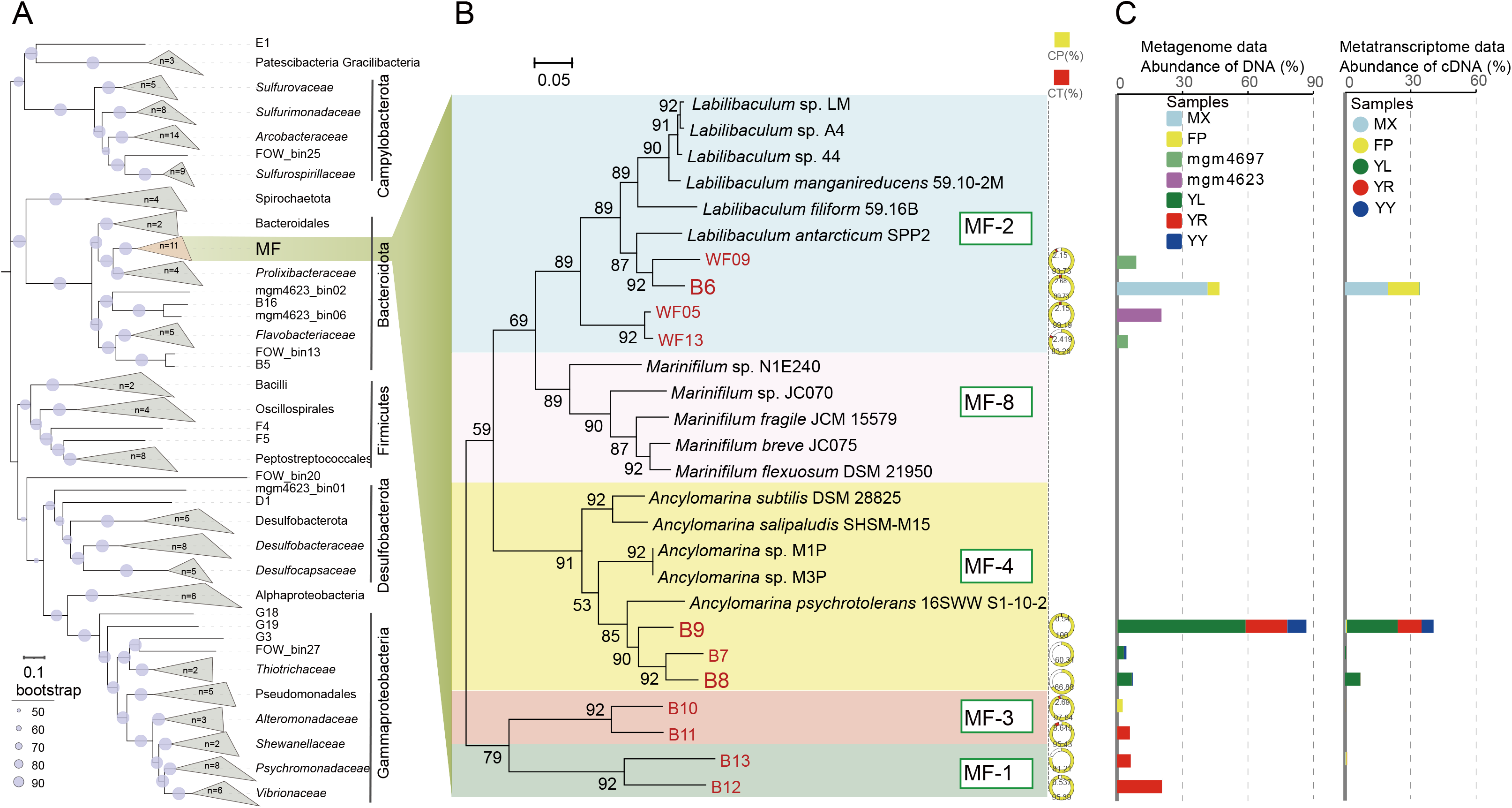
Phylogenomic tree and abundance profiles of MF taxa in the OM-enriched assemblages. (**A**) Phylogenetic tree constructed based on alignments of concatenated marker genes from the MAGs obtained in this study. The numbers in each clade represent the number of MAGs from this study. (**B**) Phylogenetic tree of the MF lineage constructed by using concatenated marker genes, including 11 MAGs obtained from this study and 16 genomes from pure cultures. CP, the completeness of MAGs. CT, the contamination of MAGs. (**C**) The abundance profiles of MF members in the OM-enriched assemblages with wood chips, wheat bran, fish scale, fish tissue, and fish oil *in situ* in the deep sea based on metagenomic (**left**) and transcriptomic (**right**) datasets. MX, wood chips; FP, wheat bran; YL, fish scale; YR, fish tissue; YY, fish oil (DHA and EPA); mgm4697 and mgm4623 are the public wood-falls.

These MF MAGs were predominant in those five microbial assemblages, exhibiting 8.4-68.5 % genomic DNA abundance (**Table S5**), as well as in the wood-falls. However, different subgroups showed a preference for organic substrates. B6, WF05, WF09, and WF13, all affiliated with MF-2, were only enriched with the plant detritus (wood chip and wheat bran), while others were enriched with the animal tissue (fish tissue and fish scales) and /or fish oil (**Fig. 2C**). This indicated that the existence of different metabolic mechanisms between different subgroups.

### MF-2 bacteria involved in the hydrolysis of polysaccharides

MF-2 bacteria could encode 527 to 1041 carbohydrate-active enzymes, which were more than that in the other three subgroups (MF-1, 3, and 4) (**Table S6**). Moreover, the genes encoding glycoside hydrolases (GHs) were significantly (*P* < 0.001) enriched in MF-2 bacteria compared to the other three subgroups (**Fig. S3**), which implied MF-2 possessed a high potential for polysaccharide hydrolysis and substrate range. For example, No GH family associated with cellulose or xylan hydrolysis was found in bacteria of MF-1, 3, and 4, but many were found in MF-2 bacteria, e.g., GH43, GH5, GH26, GH9, GH10, and GH51 (**Table S6**).

MAG B6 dominating the consortia enriched with wood chips and wheat bran harbored 263 GH genes, more than half of which were predicted to contain signal peptide sequences (**Table S7** and **Fig. S3**). These GH genes were assigned to 60 GH families and formed at least 32 of polysaccharide utilization locis (PULs) (**Fig. S4** and **Fig. S5**), which were more than that in *Polaribacter* MAGs (less than ten PULs) thriving on spring algal blooms (18). These PULs in MAG B6 were mostly involved in the hydrolysis of hemicellulose, pectin, starch, cellulose, polysaccharides containing N-acetyl groups (such as peptidoglycan and chitin), and various oligosaccharides (**Fig. S5**). Moreover, like other marine Bacteroidetes bacteria present in algae blooms (18, 39, 40), MAG B6 could encode at least 34 pairs of SusCD-like complexes (**Fig. S5**), which were found to be responsible for binding and transporting oligosaccharides into cells (41). A study on wood-fall showed that there were also many SusCD-like genes present in the metagenome data (19).

Consistent with the genomic features for saccharide metabolism, all the genes in MAG B6 involved in the hydrolysis of polysaccharides, in addition to oligosaccharide and monosaccharide metabolism, were actively transcribed in the wood chip and wheat bran consortia (**Fig. 3A**, **Table S7** and **Table S8**). In particular, 71.5 % and 30.8 % of transcripts assigned to GH genes in the wood chip and wheat bran consortia were generated from MAG B6, respectively (**Fig. 3B**). The genes involved in the hydrolysis of hemicellulose, pectin, starch, N-acetyl-containing polysaccharides, and oligosaccharides were occupied > 89 % of the total GH transcripts in MAG B6 (**Fig. S6**). The polysaccharide metabolism via MAG B6 is shown in **Figure 4A**. Above results indicated that MAG B6 contributed significantly to the degradation of plant detritus *in situ* in the deep sea.

**Figure 3.**
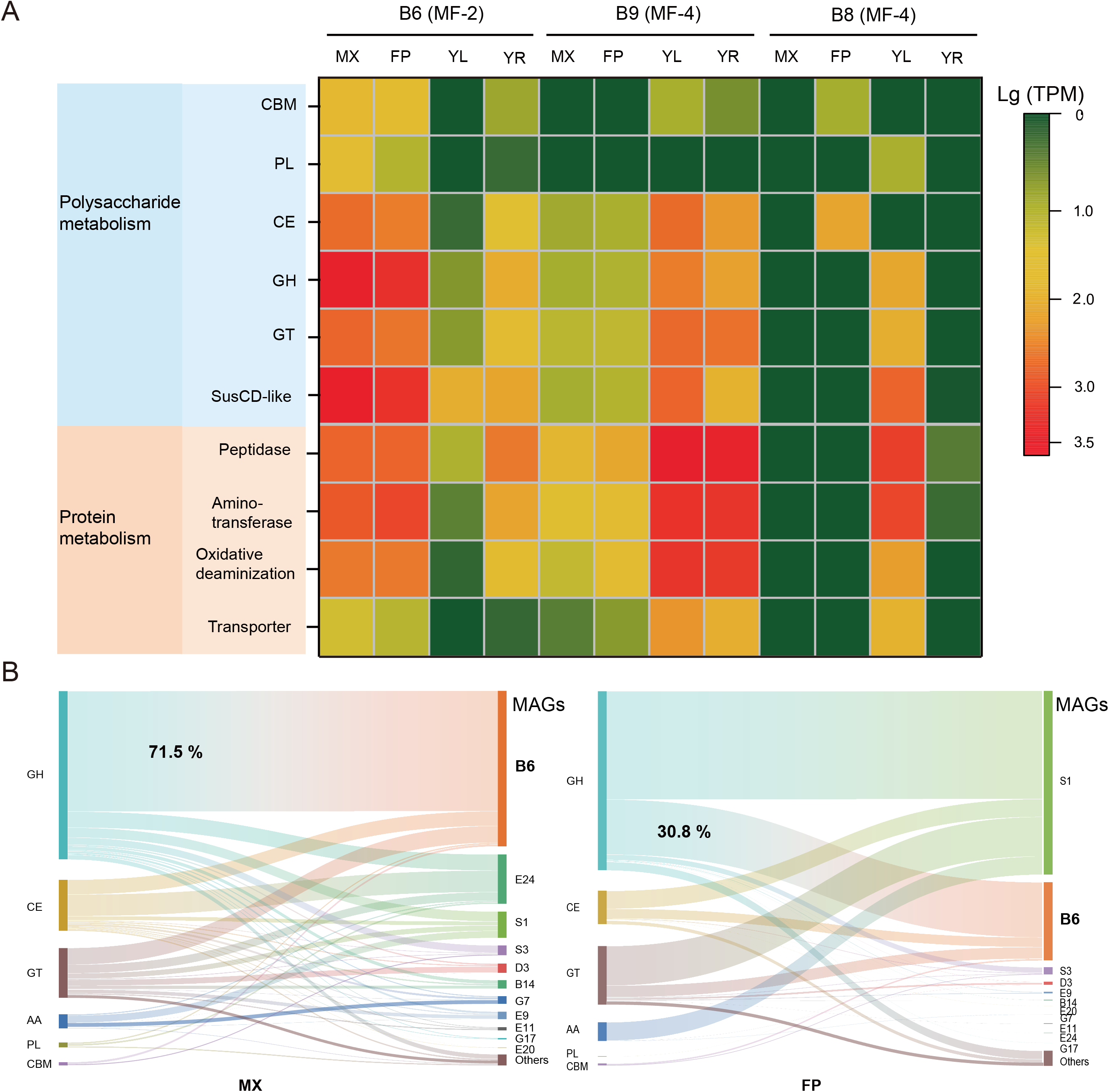
Transcriptional profiles of metabolic pathways involving polysaccharide and protein degradation. (**A**) Metabolic transcriptional profiles associated with polysaccharide and protein degradation for the dominant MF members B6, B8, and B9 in four POM-enriched assemblages with wood chips, wheat bran, fish scale, and fish tissue. (**B**) Transcriptional distribution patterns of gene sets related to polysaccharide degradation in different MAGs in MX- and FP-enriched consortia. MX, wood chips; FP, wheat bran; YL, fish scale; YR, fish tissue. GH, glycoside hydrolase; CE, carbohydrate esterase; GT, glycosyltransferase; CBM, carbohydrate-binding module; PL, polysaccharide lyase; AA, auxiliary activity.

**Figure 4.**
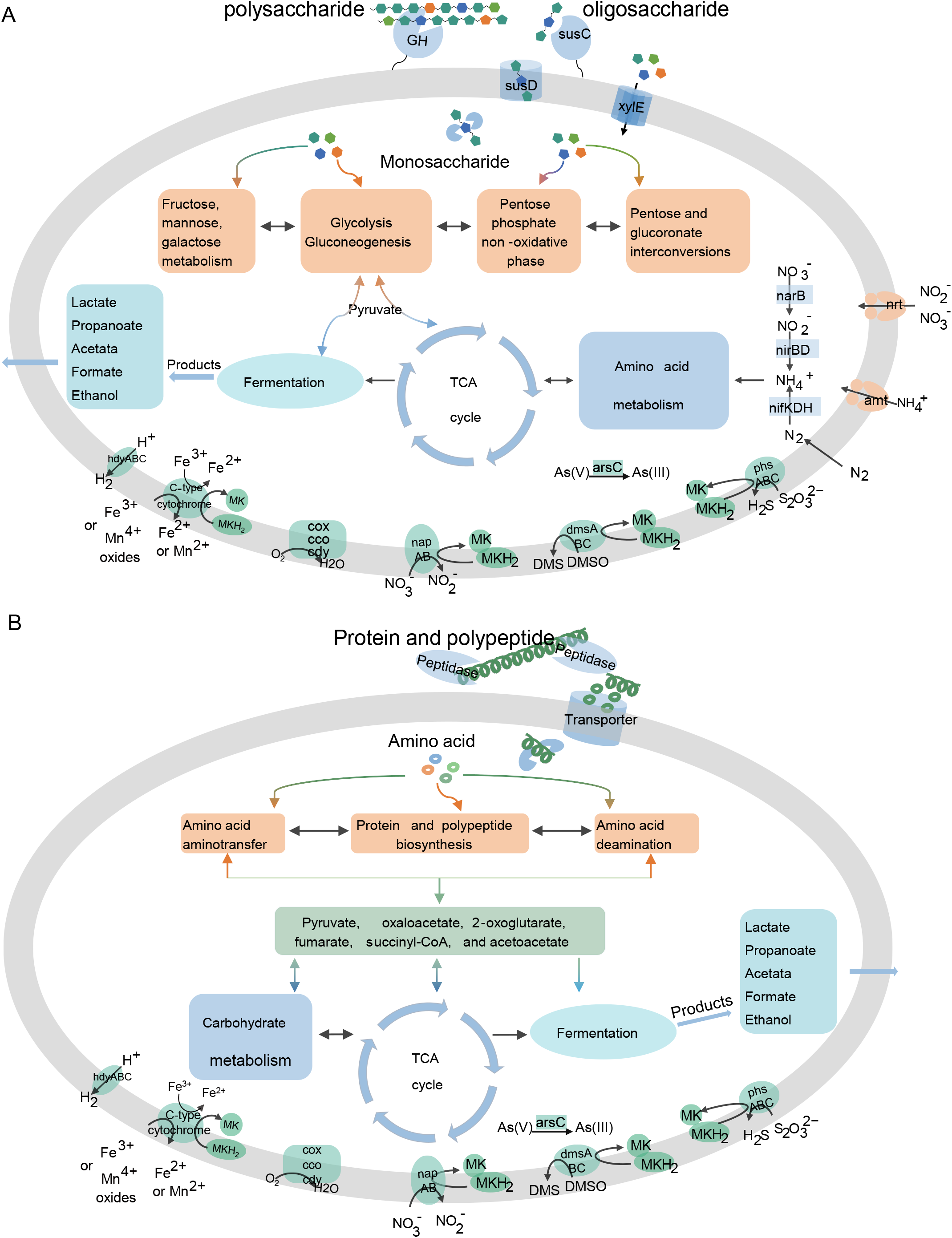
The metabolism of MF B6 and B9. (**A**) *In situ* depolymerization, degradation, and transformation of extracellular polysaccharides via MF B6 coupled with various respiration, fermentation, and other elemental cycling processes in the deep sea. (**B**) *In situ* depolymerization, degradation, and transformation of extracellular protein or polypeptide via MF B9 coupled with various respiration, fermentation and other elemental cycling processes in the deep sea. MF B6 and B9 were the dominant species in plant detritus and animal tissue enrichments, respectively. They are the keystones in the degradation of polysaccharide-POM and protein-POM *in situ* in the deep sea. Both nitrogen fixation and nitrate assimilation pathways were present in MF B6 but not in MF B9.

Among these GH genes in MAG B6, the gene (gene ID 27_12) exhibited the highest and second transcriptional activity in wheat bran and wood chip consortia, respectively (**Table S7**). This gene was closely related to the gene *xyn10b* (PDB id: 2W5F) of the bacterium *Clostridium thermocellum* (42) and encoded a CMB domain and an endo-1,4-beta-xylanase catalytic domain affiliated with the GH10 family (**Fig. S7A**). This result indicated this xylanase played a major role in the degradation of plant detritus *in situ*.

### The potential of lignin oxidation by MF-2 subgroup

Since lignin is one of the main components in wood chips, we also analyzed whether MF B6 has the potential to oxidize lignin. The genome data showed that one manganese superoxide dismutase (MnSOD) gene and one catalase-peroxidase (KatG) gene were detected in MAG B6, both of which have been confirmed to oxidize lignin and/or lignin derivatives (43, 44). MnSOD encoded by MAG B6 shared a high amino acid similarity of 64.0 % with two MnSODs from *Sphingobacterium* sp. T2, and also contained the structurally and functionally important residues (**Fig. S8**). *Sphingobacterium* sp. T2 could produce hydroxyl radical from hydrogen peroxide via its MnSODs, and then hydroxyl radical oxidizes the lignin via C*_α_*-C*_β_* and aryl-C*_α_*cleavage and demethylation (43, 45).

The upstream and downstream genes of MnSOD in MAG B6 were annotated to be iron superoxide dismutase (FeSOD) and efflux enzyme DinF, respectively (**Fig. 5A**). DinF belongs to the subfamily of multidrug and toxic compound extrusion (MATE)-like protein and is involved in the extrusion of reactive oxygen species (46). Since MnSOD from MAG B6 did not have signal peptides or encapsulin gene adjacent to it like DyP-type peroxidase from *Rhodococcus jostii* RHA1 (47), suggesting that it was probably not secreted extracellularly. Here, we constructed a possible model of lignin oxidation by MAG B6 (**Fig. 5A**). FeSOD in B6 provided hydrogen peroxide from superoxide to MnSOD; MnSOD produced hydroxyl radical via one-electron reduction of hydrogen peroxide; finally, hydroxyl radical was promptly excreted out of the cell by DinF and oxidized the lignin outside the cell (**Fig. 5A**).

**Figure 5.**
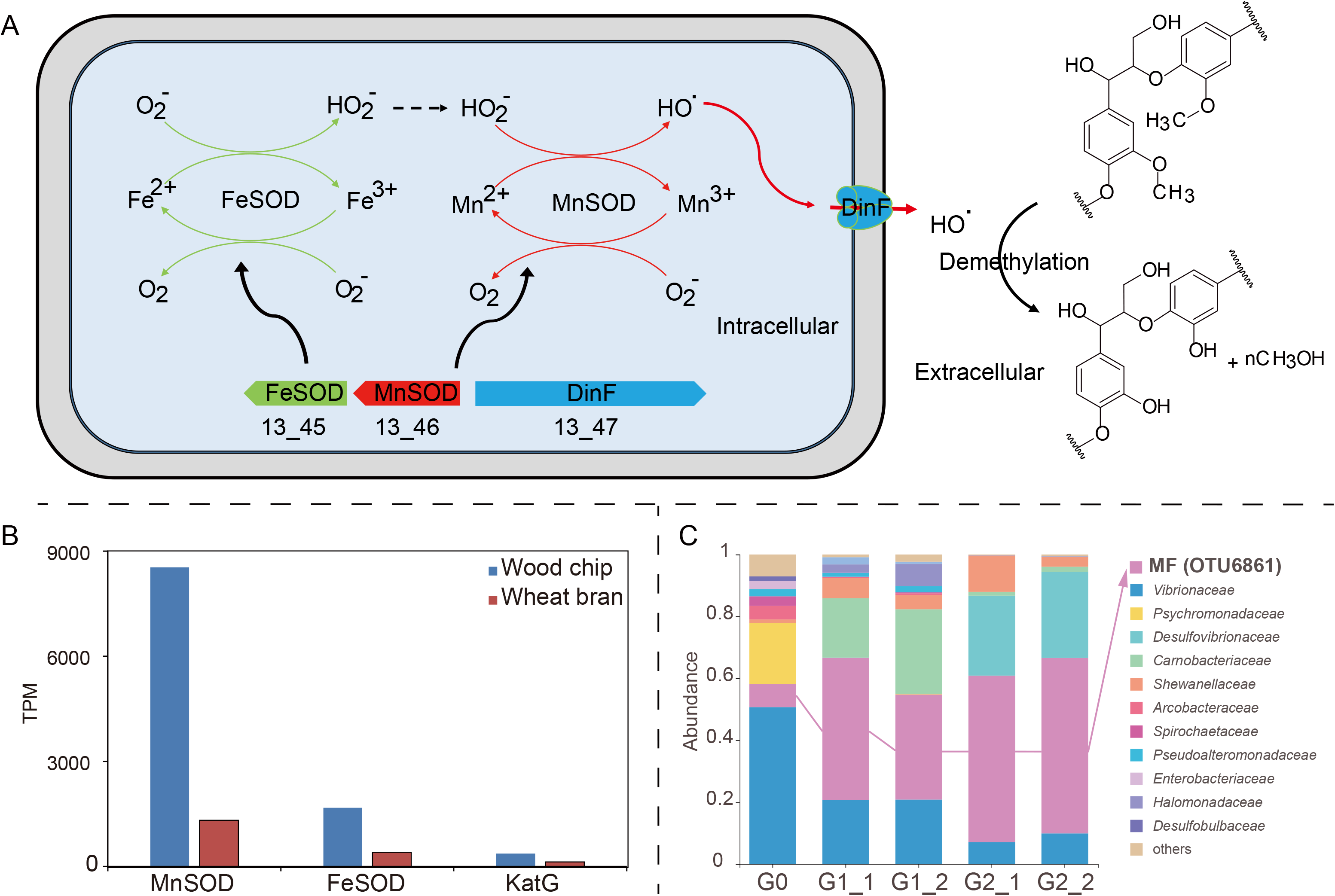
Proposed mechanism of lignin oxidation by MF bacteria and the bacterial community succession incubated with lignin. (**A**) Schematic diagram of the proposed mechanism of lignin oxidation by MF bacteria. We hypothesized that hydrogen peroxide produced by FeSOD will be provided to MnSOD, and MnSOD then reduces hydrogen peroxide to hydroxyl radicals. The hydroxyl radicals produced in the cell will promptly excreted out of the cell by the DinF and oxidize the lignin by demethylation outside the cell. (**B**) The transcriptional abundance of the genes of MAG B6 involved in lignin oxidation in wood chips and wheat bran enrichments. MnSOD and FeSOD showed higher transcriptional activity in wood chip enrichment than that in wheat bran enrichment. (**C**) Succession of bacterial community incubated with lignin as the sole organic carbon and energy. The details are shown in the Supplementary material.

In addition, this cluster including MnSOD, FeSOD, and DinF was also detected in other MF-2 bacteria, while it was not found in most MAGs belonging to MF-1, MF-3, or MF-4 (except MAG B9).

Metatranscriptomic analysis showed that genes encoding MnSOD and FeSOD were notably transcribed at a higher level in wood chip enrichment than that in wheat bran enrichment (**Fig. 5B**), which implied MAG B6 may oxidize lignin or its derivatives *in situ* in the deep sea. Furthermore, the community succession with lignin as the sole carbon and energy source at the laboratory, showed the relative abundance of OTU6861 that shared the same 16S rRNA gene sequence as MAG B6, increased significantly with incubation time (0, 0.5, and 1 year) (**Fig. 5C** and details were shown in **Supplementary material**), which further suggested that it may oxidize and utilize lignin to grow.

In the wood-fall ecosystem, it is regarded that the degradation of wood polysaccharides is mainly carried out by macrofaunal borers such as *Xylophaga* spp. (38, 48). Researchers believed that bacteria, including the MF clade, use only small organic compounds, such as sucrose, directly derived from wood or excretions from these macrofaunal borers *in situ* (29, 38). Therefore, the oxidation of lignin in the wood-fall and the keystone bacteria involved in this process remain mysterious. In this study, we gained insight into the ecological role of MF-2 bacteria in the degradation of polysaccharides and oxidation of lignin *in situ* in the deep sea. Furthermore, based on the above results, we believe that MF members present in water columns are possibly involved in the depolymerization of POM, since carbohydrates are known as one of the main components of POM in marine water columns (49) and the extracellular enzyme activity of glycoside hydrolases is abundantly detected in POM (14). Additionally, MF bacteria are present in oligotrophic sediment (35), which may be sustained by decomposing and utilizing those buried refractory DOM derived from POM and DOM by the microbial carbon pump (5) and buried refractory humic acid materials, as well as refractory components (such as peptidoglycan) derived from the cell walls of dead microbes.

### Protein hydrolysis and utilization by subgroups of MF-1, −3, and −4

Bacteria of the MF-1, −3, and −4 subgroups dominated the communities of protein enrichments *in situ*, especially the two bacteria of MF B9 and MF B8, both of which belong to subgroup MF-4 (**Fig. 2C**). There were 119 and 82 peptidase-encoding genes retrieved from MAG B9 and B8, respectively (**Tables S9- S11**). Notably, approximately half of the peptidases encoded by MF B8 and B9 possess the N-terminal signal peptide (**Tables S10-S11**), indicating that they are secreted extracellularly and responsible for the degradation of extracellular protein and/or peptide (50). Moreover, some genes encoding aminotransferases and oxidative deaminases involved in amino acid metabolism were also detected, as well as central amino acid metabolism or degradation pathways, such as those for alanine, aspartate, and glutamate (**Fig. 4B** and **Table S12**).

Furthermore, metatranscriptomic datasets from two enrichments with fish scale and fish tissue showed that MAGs B9 and B8 exhibited high transcriptional activities of functional genes responsible for polypeptide degradation, in addition to amino acid metabolism (**Fig. 3A**). Particularly, 78.8 % and 48.3 % of transcripts in the two consortia assigned to these secreted peptidases were generated from MAG B9 and B8, respectively, indicating that both bacteria contributed significantly for the degradation of polypeptides *in situ* in the deep sea.

### Differentiation among MF bacterial subgroups in nitrogen fixation and nitrogen source acquisition

Nine members of ten within subgroup MF-2 harboured a complete nitrogen fixation gene cluster (*nifHDKEBV*), along with related transcriptional regulatory genes (**Fig. 6B**). In contrast, the MF-1, MF-3, and MF-4 bacteria dominating the animal tissue-enrichments did not encode the abovementioned nitrogenase (**Table 1** and **Table S12**). The enzymes of NifHDK encoded by MF-2 bacteria contain the important conserved residues (51), like other nitrogen-fixing bacteria (**Fig. 6C**). These NifHs shared relatively high amino acid similarities (67-72 %) with nitrogen-fixing bacteria *Clostridium pasteurianum*, *Azotobacter vinelandii*, and *Klebsiella pneumoniae* (52) (**Table S13**). Moreover, we observed nitrogen fixation in the laboratory by an MF-2 isolate (*Labilibaculum* sp. LM in **Figure 2**) of mangrove sediment origin (data not shown).

**Figure 6.**
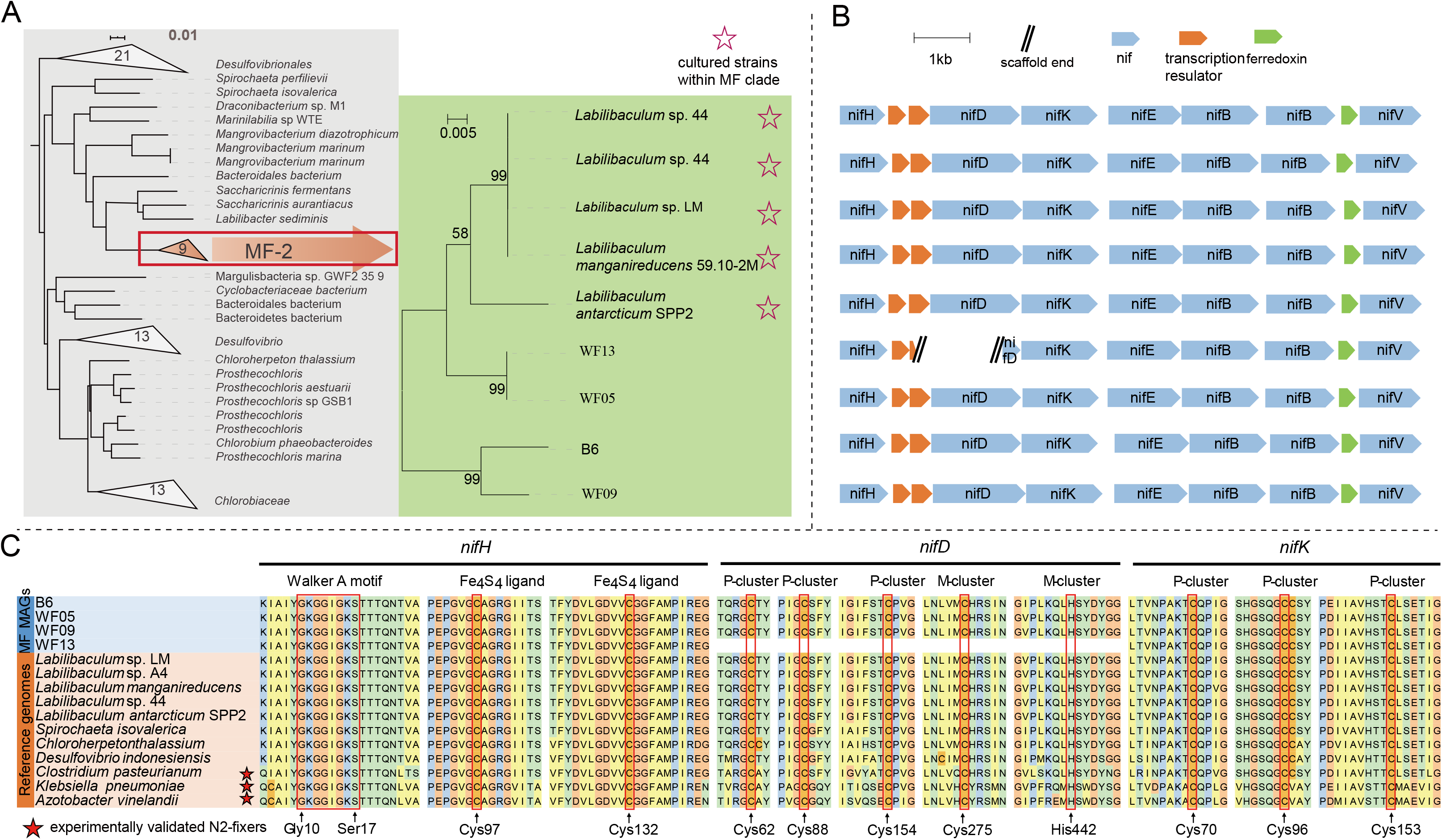
Phylogenetic and genomic analyses of nitrogenase. (**A**) Phylogenetic tree of *nifH* of MF-2, together with other closely related *nifH* genes from the NCBI database. (**B**) Schematic diagram of the nitrogenase gene cluster of the MF-2 MAGs from this study and reference strains. (**C**) Conserved amino acids for the key residues in three important proteins (*nifHDK*).

In addition to nitrogen fixation gene, a gene cluster for assimilatory nitrate reduction (*narB*, *nirBD*, and *nrtABCD*) was detected only in MAG B6, but not in the genomes of MF-1, −3, and −4 subgroup bacteria (**Table S12**). The metatranscriptomic data revealed these genes from MAG B6 involved in nitrogen fixation and assimilatory nitrate reduction were transcribed during polysaccharide/lignin degradation, although their transcripts were low compared to some GH genes (**Table S12**). These results indicated that the pattern of nitrogen source acquisition may be one of the key metabolic features determining MF bacterial specialization and adaptation to different types of OM and that MF-2 members may play an important role in balancing the disproportion of nitrogen and carbon sources, especially when the available nitrogen sources are scarce and polysaccharide or lignin is saved as growth substrates.

This is the first report that *Marinifilaceae* bacteria within the phylum Bacteroidetes have nitrogen-fixing potential. A previous report showed that abundant nitrogen-fixing genes are derived from Bacteroidetes in the termite gut (53, 54), suggesting that Bacteroidetes may play a significant role in the nitrogen balance of the intestinal flora. Similarity, MF members are also detected in the gut of marine animals in addition to the surface of marine plants and animals, e.g., barnacles and corals (**Fig. S2** and **Table S3**) (55–57). Those MF bacteria may act as probiotics to provide ammonia nitrogen to the host by nitrogen fixation *in situ*. The wide distribution of MF-2 in the ocean indicates the universality of nitrogen fixation in the marine environment, as recently reported regarding the diversity of diazotrophs in deep-sea sediment (58).

### Versatile anaerobic respiration mechanisms by MF bacteria

In addition to the microaerophilic respiration (*coxABC*, *ccoNP*, and *cydAB*), most MF bacteria possessed the potential to use thiosulfate, dimethyl sulfoxide (DMSO), and arsenate as the terminal electron acceptors but were incapable of sulfate reduction (**Fig. 7AB** and **Table S14**). In addition, both *L*. *manganireducens* 59.10-2M^T^ and *L*. *filiforme* 59.16B^T^ have been confirmed to reduce Fe^3+^ and Mn^4+^ ions and/or oxides during glucose fermentation, presumably through a cytochrome c gene (35). This gene was also detected in most MAGs (**Fig. 7AB** and **Table S14**). At present, the metal-reducing (Fe^3+^ and Mn^4+^) bacteria mainly belong to Firmicutes, Proteobacteria, Deferribacteres, and Deinococcus-thermus (59, 60). Moreover, MAG B6 and WF09 additionally possessed a gene cluster for dissimilatory nitrate reduction (*napABCDFGH*) (**Fig. 4** and **Fig. 7AB**).

Fermentation was additionally found to be a common strategy among MF members, even in the presence of some alternative terminal electron acceptors. Most MF bacteria in this study had the potential to generate formate (*pflD*), ethanol (*aldH* and *adhE*), acetate (*acdAB* and *ackA*), propionate (*mmdAC* and *pccB*), and lactate (*lldF* and *dlD*) via fermentation (**Fig. 7AB** and **Table S14**), like *L*. *manganireducens* 59.10-2M^T^ and *L*. *filiforme* 59.16B^T^ (35). In addition, almost all members within MF-2 and MF-4 possess three groups of [FeFe]-hydrogenases (Group A3, B, and C1; *hydABC*, *hydS*, and *hydM*) (**Fig. 7AB**, **Fig. S9-S10**, and **Table S14**), all of which are typically involved in strictly anaerobic fermentation for hydrogen production (61). These fermentation products can serve as carbon sources for sulphate-reducing bacteria, e.g., *Desulfobulbaceae* and *Desulfobacteraceae,* also dominating the animal tissue- and plant-detritus enrichments (33).

Metatranscriptomics analysis showed that all the genes involved in anaerobic respiration and fermentation were actively transcribed in MAG B6, MAG B8, and MAG B9 during corresponding OM degradation (**Fig. 7B** and **Table S14**). Particularly, the genes encoding putative Fe^3+^ and Mn^4+^ ions and/or oxides of reductase exhibited obviously high transcriptional levels regardless of the organic substrates, and the gene encoding (*mmdAC* and *pccB*) for propionate fermentation showed significantly high expression levels (**Fig. 7B**), which suggested MF members may prefer Fe^3+^ and Mn^4+^ ions and/or oxides as electron acceptors and ferment OM to generate propionate during OM degradation *in situ* in the deep sea. The energy gained from Fe^3+^ and/or Mn^4+^ reduction is high, comparable to that from oxygen as electron acceptor. The above results indicated that MF bacteria used various substances as terminal electron acceptors coupled with diverse fermentation types during the degradation of various OMs *in situ* in the deep sea.

**Figure 7.**
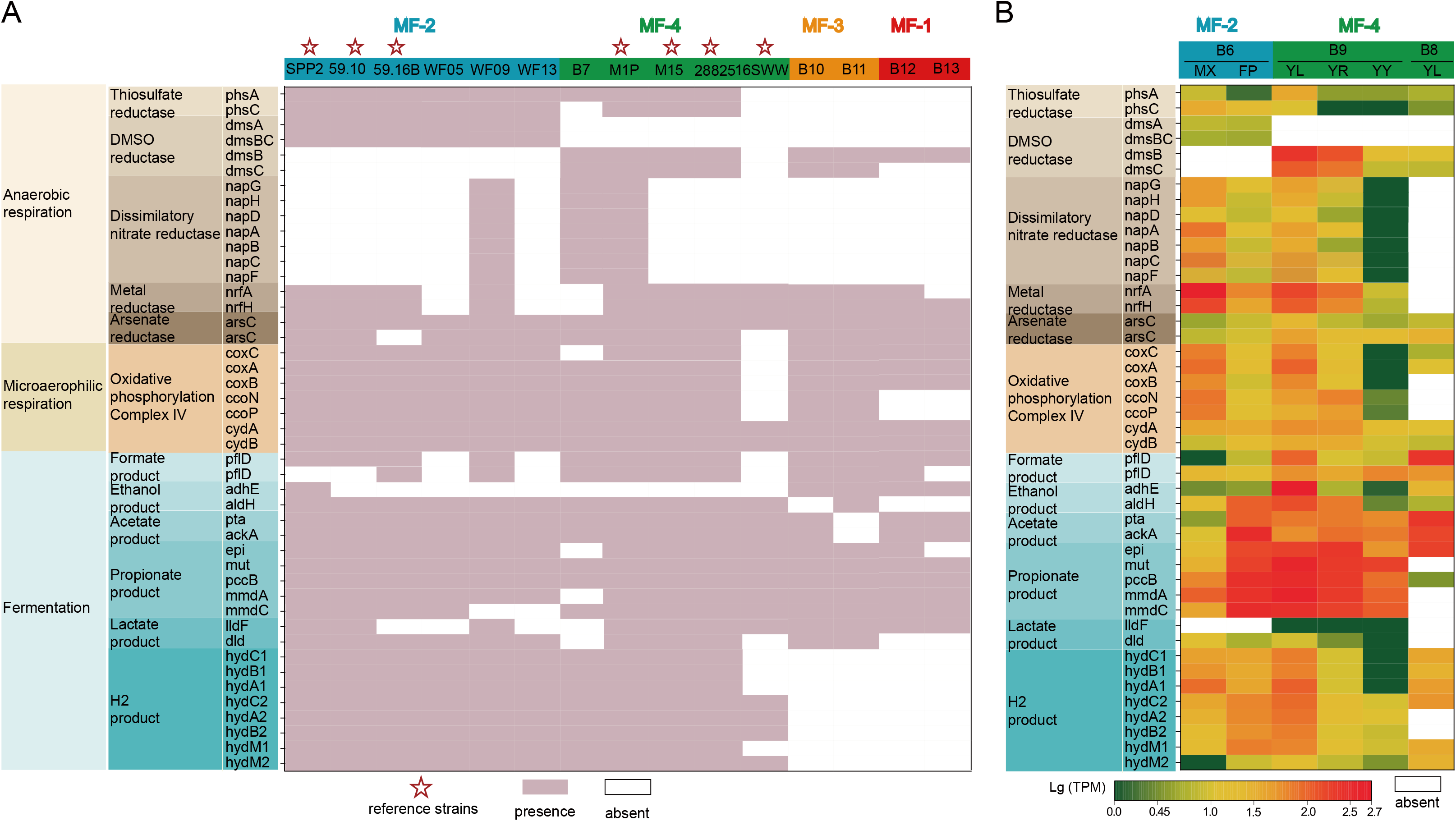
Cell respiration and fermentation pattern in MF members. (**A**) The genes present or absent in the genomes of MF members involved in cell respiration and fermentation. (**B**) Transcriptomic profiles of related genes in MAG B6, B8, and B9. SPP2, *L*. *antarcticum* SPP2; 59.10, *L*. *manganireducens* 59.10-2M; 59.16B, *L*. *filiform* 59.16B; M1P, *A*. sp. M1P; M15, *A*. *salipaludis* SHSM-M15; 28825, *A*. *subtilis* DSM 28825; 16SWW, *A*. *psychrotolerans* 16SWW S1-10-2; B6-B13, WF05, WF09, and WF13 were obtained from this study.

Biodegradation of OM will inevitably result in oxygen depletion, which not only occurred in the past but also occurs currently in global oceans, e.g., oxygen-minimum zones (62) and wood-fall (38). Similarly, the globally enlarging anoxic areas are the consequence of abundant OM input from rivers and eutrophication in estuarine areas (63). Moreover, fermentation is often found in other OM-rich environments, such as cold seep (64) and petroleum seep in deep-sea sediments (65).

## CONCLUSION

This study highlighted the ecological roles of the bacteria of the family *Marinifilaceae* within the phylum Bacteroidetes in the depolymerization, degradation, and transformation of different types of macromolecular OM in global oceans, as summarized in the diagram **Figure 8**. Their diversity and wide distribution in marine environments, especially in the chemoheterotrophic ecosystems represented by wood falls and whale falls, strengthen the findings of our *in situ* incubation experiments. These bacteria diverge not only in phylogeny but also in metabolism for the degradation of different macromolecules. It is noteworthy that MF-2 subgroup bacteria possess overwhelming advantages in polysaccharide hydrolysis and potential lignin oxidation, and intriguingly, they also possess nitrogen fixation potential, which distinguishes them from the predominant members in animal tissue- and fatty acid-supported consortia. These MF bacteria dominate the macromolecule-decomposing bacterial assemblages, all conduct fermentation and can respire diverse electron receptors and relay with sulfate reducers via fermentation products. Therefore, MF members are key players in OM mineralization in both anoxic and oxic marine environments and are coupled with sulfur, nitrogen, and metal elemental biogeochemical cycles.

**Figure 8.**
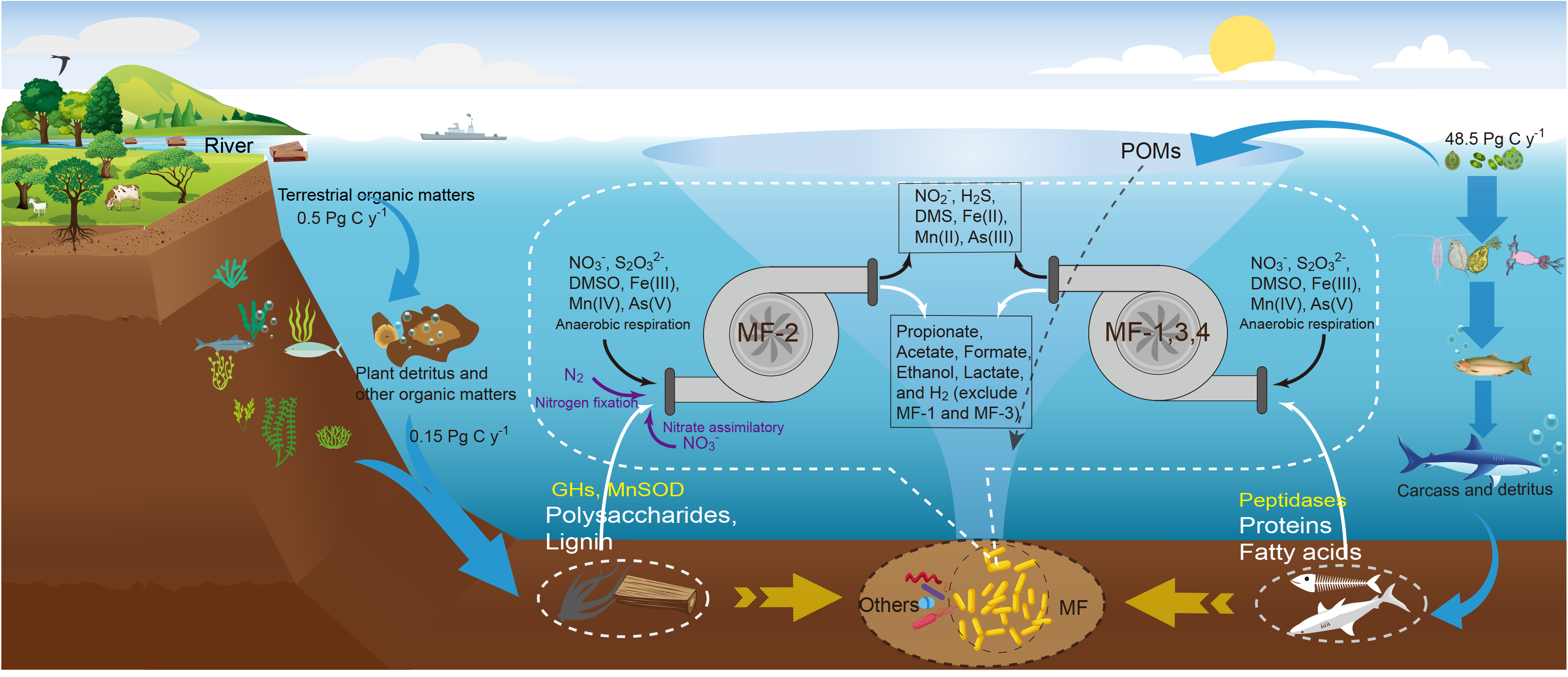
Schematic depiction of the ecological role of MF taxa in the global oceans. From this study, we gained insight into the ecological role of MF bacteria in the depolymerization of POM derived from animal or plant detritus *in situ* in the deep sea under anoxic conditions, simultaneously coupled with nitrogen, sulfur, and metal elemental biogeochemical cycling. MF members are the predominant group in the assemblages thriving on newly input plant detritus and animal tissue *in situ* in the deep sea. In particular, MF-2 members prefer to biodegrade polysaccharides and lignin, while MF-1, MF-3, and MF-4 prefer to biodegrade proteins or polypeptides. Omics data show that MF-2 members have significantly more glucoside hydrolase-associated genes than the other three groups (MF-1, MF-3, and MF-4), possessing overwhelming advantages in polysaccharide hydrolysis. Additionally, both nitrogen fixation (present in almost all MF-2 members) and the nitrate assimilation pathway (only in B6 of MF-2) likely provide a strong survival advantage for MF-2 members, especially when the available nitrogen sources are scarce and polysaccharides and lignin are saved for growth substrates. On the other hand, no nitrogen fixation or nitrate assimilation pathway was detected in the other three MF subgroups thriving on animal tissue; therefore, they are likely specialized in degrading proteins or polypeptides with many peptidases.

## MATERIALS AND METHODS

### Incubation samples

Previously, we obtained a large number of plant detritus- and animal-incubations from the deep sea of the marginal sea (the South China Sea) and ocean (the Pacific Ocean and the Indian Ocean) respectively using a deep-sea *in situ* microbial incubators. In this study, we selected five representative incubation samples to further analyze the metabolic potential of MF subgroup in the *in situ* environment through metagenomic and macrotranscriptomic data. The five incubations respectively emended with wood chip, wheat bran, fish scale, fish tissue, and fish oil (DHA and EPA) were incubated at a flat-topped seamount in the Pacific Ocean (20.4059567° N, 160.7700883° E; 1622 m water depths) for 348 days. The detailed enrichment process and organic substrates refer to this previous article (33).

### DNA and RNA extraction

The enriched biomass in the liquid phase was filtered on a 0.22-μm pore size polycarbonate membrane (Millipore, USA) and a half was used for DNA extraction and the other half for RNA extraction. Total DNA was extracted and purified with a DNeasy PowerWater Kit (Qiagen, Germany) according to the manufacturer’s protocol. Total RNA was extracted using the RNeasy Mini Kit (Qiagen, Germany) according to the manufacturer’s instructions. The concentration and quality of the extracted DNA and RNA were determined using a NanoDrop2000 spectrophotometer (Thermo Scientific) and gel electrophoresis, respectively. All DNA and RNA samples were stored at −80 °C until further processing.

### Cloning of 16S rRNA gene and sequencing

The extracted DNA was amplified using the broad-range bacterial-specific 16S rRNA gene primer pair 27F: 5’-AGAGTTTGATCCTGGCTCAG-3’ and 1492R: 5’-ACGGCTACCTTGTTACGACT-3’. Amplicons were purified by using the QIAquick PCR Purification Kit (Qiagen Inc., Valencia, CA) and clone libraries were constructed with the TOPO TA cloning kit (Invitrogen) according to the manufacturer’s instructions. Inserts from the clones were amplified with the M13F/R primers and Sanger sequenced from both ends by the Majorbio Company (Shanghai, China).

### Global distribution pattern

To reveal the global distribution of the enriched MF species in the deep-sea enrichments, we collected the microbial data containing MF reads by using the Integrated Microbial Next Generation Sequencing (IMNGS, https://www.imngs.org, last accessed on May 5, 2019) server, which pigeonholes all available raw sequence read archives retrieved from the International Nucleotide Sequence Database (GenBank, DDBJ, and EMBL) and allows users to conduct comprehensive searches of SSU rRNA gene sequences. In this study, representative sequences (the nearly full-length 16S rRNA gene) were subjected to a similarity search against the IMNGS database using a similarity threshold of 95 % and a min length of 200 bp. A total of 1026 samples were retrieved and their metadata including source, latitude, longitude, and depth of seawater was obtained from NCBI (**Table S1**).

### Metagenomic sequencing, assembly, and binning

Metagenomic library preparation and DNA sequencing using Illumina NovaSeq 6000 platform (PE 150-bp mode) were conducted at the Majorbio Company (Shanghai, China). Raw metagenomic reads were QC-processed by using the fastp v0.19.3 with default parameters (66). Possible animal or plant reads from substrates were removed by mapping to the genomes of *Pinus*, *Triticum aestivum*, and *Larimichthys crocea* using Bowtie2 (identity ≥ 0.9) (67). Filtered reads were individually assembled *de novo* by MetaSPAdes v3.13.0 with the settings “-k 21,33,55” (68). Previous studies on wood-falls showed the presence of MF members (19, 38), so we additionally retrieved metagenomic data from the MG-RAST database (https://www.mg-rast.org/). Their accession numbers are mgs476148, mgs476151, mgs476154, mgm4623131.3, mgm4623132.3, and mgm4623133.3. These data were QC-processed and co-assembled using the same method.

The binning process was performed by using the Metawrap pipeline v1.2.1 based on three methods of metabat2, maxbin2, and concoct (69). Two modules of “bin_refinement” and “reassemble_bins” both in the Metawrap pipeline were then implemented with the parameters “-c 50, -x 10” (69). Afterward, MAGs with the thresholds of > 50 % completeness and < 10 % contamination were picked for subsequent analysis. All binning results were combined and dereplicated using dRep (-comp 40 -con 15 --run_tax -g options). After dereplication, a total of 110 dereplicated MAGs were obtained from our five deep-sea *in situ* incubations and 35 MAGs were obtained from wood-falls. The completeness and contamination of each MAG were estimated by CheckM v1.0.12 (70). The coverage of each MAG in each sample was calculated by using Salmon software in the Metawrap pipeline with the quant_bins module (69).

### Taxonomic assignments and ANI calculation

Taxonomic assessment of each MAG was performed using GTDB-Tk with the database of R04-RS89 (71). The ANI values were calculated with FastANI v1.1 (72).

### Functional annotations

Protein-coding genes of each MAG were predicted using Prodigal v2.6.3 (73). Protein sequences were functionally annotated against databases, including KEGG (74), eggNOG (75), Pfam (76), MEROPS (77), and CAZy (78). The online software KAAS v2.1 (74) (https://www.genome.jp/kegg/kaas/) was applied for homology searching against the KEGG database with the GHOSTZ program. Eggnog-mapper v1.0.3 software was used to annotate with the Diamond Blastp (v0.8.36.98) method (75). To identify carbohydrate degradation-related enzymes, we used the online dbCAN2 meta server (http://bcb.unl.edu/dbCAN2/) by HMMER against the CAZy database (78). Pfam 31.0 (76) was used as the reference database for the annotation of peptidases, aminotransferases, and transporters of oligopeptides and amino acids by HMMER v3.1b2 (cut-offs: e value: 1e-10, best hits reserved). Additionally, the MEROPS database (Release 12.1) (77) was used for peptidase annotation by Diamond Blastp v0.8.36.98 (cut-off: e value: 1e-10, best hits reserved) (79). When a protein sequence was annotated to the same peptidase family with the above two databases, its annotation result was accepted and then used for subsequent analyses. The prediction of signal peptides was carried out by using the online SignalP-5.0 Server (http://www.cbs.dtu.dk/services/SignalP/). The three-dimensional structures of certain key encoding proteins were predicted by using the online Phyre2 web portal (80).

### Metatranscriptomic sequencing and mapping

Metatranscriptomic library preparation and cDNA sequencing using Illumina NovaSeq 6000 platform (PE 150-bp mode) were conducted at the MAGIGENE Company (Guangzhou, China). Raw metatranscriptomic reads were QC-processed by using the fastp v0.19.3 with default parameters (66), and the resulting clean reads without rRNA were mapped to each MAG using Bowtie2 with default parameters (67). Afterward, the counting of fragments (PE reads) assigned to each gene in MAG was carried out using the FeatureCounts program with the parameters “-p -F GTF -g ID -t CDS -s 1 -M --fraction” (81). The transcript of each gene in each MAG was normalized with the TPM (transcripts per million) (82).

### Phylogenetic analysis

16S rRNA gene homology searches within GenBank were performed using blastn. A total of 338 sequences belonging to MF family were retrieved (**Table S2**). Multiple sequence alignment of 16S rRNA gene amplicon was performed using the MAFFT program with default parameters, and a phylogenetic tree was reconstructed with the neighbour-joining method (https://mafft.cbrc.jp/alignment/server/). For the phylogenetic tree of hydrogenase, a neighbour-joining tree was constructed by using MEGA v7.0 software (83). The reference hydrogenase catalytic subunits were retrieved from a previous study (61). For draft MAGs, the phylogenomic tree was built by the up-to-date bacterial core gene (UBCG) method (84). The reference genomes were retrieved from the EzBioCloud database (https://www.ezbiocloud.net/) in May 2021.

### Data availability

The raw metagenomic sequences, raw metatranscriptome sequences, and the MAGs have been deposited in NODE (https://www.biosino.org/node/) at the accession number OEX011321, OEX011323, and OEZ007098, respectively. The nearly full-length 16S rRNA sequences belonging to MF family acquired in this study have been deposited in GenBank under the accession numbers ON228496-ON228518.

## ACKNOWLEDGMENTS

This work was financially supported by the following projects dedicated to Z.Z.S.: the National Natural Science Foundation of China (No.42030412), the China Ocean Mineral Resources R&D Association (COMRA) program (No. DY135-B2-01), High-Tech Research and Development Program of China (No. 2012AA092102), and the Scientific Research Foundation of Third Institute of Oceanography, MNR (2019021). We thank Dr. Xiuran Yin (Faculty of Biology/Chemistry, University of Bremen, Bremen, Germany) for comments on an earlier version of the manuscript.

Z.Z.S. conceived this study’s conception, applied the financial supports, organized and performed *in situ* incubation. Z.Z.S., J.Y.L., C.M.D., and Q.L. L participated the cruises. J.Y.L. contributed substantially to the data acquisition and analysis, including metagenome and metatranscriptome. Z.Z.S., J.Y.L. and C.M.D. interpreted the results and wrote the manuscript. G.Y.W. reviewed the paper. All authors approved the final manuscript.

We declare no competing interests.

## Supplementary Figure legends

**Figure S1 Phylogenetic tree based on the 25 clone sequences belonging to MF obtained in this study and the 16S rRNA gene sequence similarities within and between the groups.** A total of 25 near-full-length 16S rRNA gene sequences affiliated with the MF clade were cloned from ten representative enrichments. The lowest similarity among subgroups was 91.2 % between MF-4 and MF-2, so we used the full-length 16S rRNA gene sequences of the four phylogenetic representatives as the query to retrieve the homologues from the NCBI database using Blastn. A total of 340 16S rRNA gene sequences were recruited under a 91.2 % similarity threshold.

**Figure S2** T**h**e **global distribution pattern of MF-1 to MF-4.** The MF bacteria thrived on plant detritus and animal tissue delivered via an *in situ* incubator are affiliated with the MF-1, MF-2, MF-3, and MF-4 groups. This result shows that they were widely distributed in marine environments, including seawater columns, marine sediments, marine animal and plant surfaces or animal intestines, and other organic-rich marine habitats, such as whale-falls, wood-falls, seagrass detritus, cold seep, and petroleum-contaminated sediment, but not found in terrestrial and freshwater environments. The data are from this study and the public database (IMNGS) and are shown in Supplementary Table 1.

**Figure S3 The number of total GH genes and GH genes containing signal peptides in each genome (A) and the comparative analysis of GH numbers (B) among the five MF subgroups.** This result indicates that MF-2 members have significantly more GH-encoding genes than the other four subgroups (MF-1, 3, 4, and 8).

**Figure S4** T**h**e **gene numbers of different GH families in MAG B6.** MAG B6 of MF-2 is the predominant species group in the polysaccharide assemblages.

**Figure S5 Polysaccharide utilization loci (PULs) in MAG B6. SusCD-like genes cluster with various GH, CBM, GT, and/or CE genes to form PULs.** There are at least 32 PULs present in MAG B6. The first gene ID in each PUL is shown on the left and hydrolytic substrate for each PUL is shown on the right. The genes marked in red are SusCD-like complex genes; the genes marked in blue are GH, CBM, GT, and/or CE genes; and the genes marked grey are of unknown function.

**Figure S6 Bar chart showing the relative transcriptional levels of different GHs in MAG B6 involved in polysaccharide and oligosaccharide hydrolysis in wood chip (MX) and wheat bran (FP) enrichments.** Those GHs involved in the hydrolysis of hemicellulose, pectin, starch, N-acetyl-containing polysaccharide, and oligosaccharide are transcribed at high levels and account for > 89 % of the total GH transcripts in MAG B6.

**Figure S7 The protein structure of key enzymes.** The three-dimensional protein structure of gene 27_12 in MAG B6 involved in polysaccharide hydrolysis (A) and the key gene 16_53 in MAG B9 involved in protein hydrolysis (B), both predicted by using Phyre2 (http://www.sbg.bio.ic.ac.uk/phyre2/html/page.cgi?id=index).

**Figure S8 The conserved amino acids for the key residues in MnSOD.** MnSODs of MF members contain the structurally and functionally important residues like MnSOD1 and MnSOD2 both from *Sphingobacterium* sp. T2; Mn(II) ligands His26, His76, Asp163, and His167; gateway residue Tyr34 and catalytic Gln144; other active site residues His30, Asn75, Trp125, Trp165, Glu166, and Tyr170.

**Figure S9** Neighbour-joining tree of amino acid sequences of the group A3 [FeFe]-hydrogenase catalytic subunit, a marker for hydrogen production during fermentative and respiratory processes. The tree shows sequences of MF members from metagenome-assembled genomes obtained in this study and cultured strains (red) alongside representative reference sequences (black). The representative reference sequences were retrieved from a previous study.

**Figure S10** Neighbour-joining tree of amino acid sequences of the group B and C1 [FeFe]-hydrogenase catalytic subunit, a marker for hydrogen production during fermentative and respiratory processes. The tree shows sequences of MF members from metagenome-assembled genomes obtained in this study and cultured strains (red) alongside representative reference sequences (black). The representative reference sequences were retrieved from a previous study.

## Supplementary Table legends

**Table S1** The metadata of 1026 samples that contained MF reads retrieved from the IMNGS and NCBI databases.

**Table S2** Newick tree file for the neighbour-joining phylogenetic tree of the MF 16S rRNA genes in Figure 1B and the 16S rRNA gene sequences used here. All sequences except those from this study were retrieved from the NCBI database.

**Table S3** Statistics of metagenome assemblies.

**Table S4** Average nucleotide identity (ANI) values among the MF genomes of MAGs and cultured strains. The data were computed with fastANI.

**Table S5** The characteristics and relative abundance of the MAGs obtained in this study.

**Table S6** The number of carbohydrate-active enzymes in each genome within the MF clade.

**Table S7** The detailed list of carbohydrate-active enzymes (CAZymes), including carbohydrate esterases (CEs), glycoside hydrolases (GHs), polysaccharide lyases (PLs), carbohydrate-binding modules (CBMs) and glycosyltransferases (GTs), detected in MAG B6, as well as the transcriptional TPM values of each gene in MX and FP enrichment.

**Table S8** Summary of various monosaccharide metabolism and TCA cycle genes detected in MF MAGs obtained in this study, as well as the transcriptional TPM values of each gene in the five enrichments.

**Table S9** The number of genes encoding peptides in each member within the MF clade.

**Table S10** The detailed list of peptidases and aminotransferases detected in MAG B9, as well as the transcriptional TPM values of each gene in the YL and YR enrichments.

**Table S11** The detailed list of peptidases and aminotransferases detected in MAG B8, as well as the transcriptional TPM values of each gene in the YL enrichment.

**Table S12** Summary of nitrogen metabolism, sulfur metabolism, various amino acid metabolism, and fatty acid degradation genes detected in MF MAGs obtained in this study, as well as the transcriptional TPM values of each gene in the five enrichments.

**Table S13** The amino acid similarities of nifH between MF-2 members and other experimentally validated N2 fixers Clostridium pasteurianum, Azotobacter vinelandii, and Klebsiella pneumoniae.

**Table S14** Summary of cell respiration-associated metabolic processes, including aerobic respiration,

## REFERENCES

1. Arndt S, et al. (2013) Quantifying the degradation of organic matter in marine sediments: A review and synthesis. Earth-Sci. Rev. 123:53–86.

2. Jørgensen BB & Boetius A (2007) Feast and famine--microbial life in the deep-sea bed. Nat. Rev. Microbiol. 5(10):770–781.

3. Orcutt BN, Sylvan JB, Knab NJ, & Edwards KJ (2011) Microbial Ecology of the Dark Ocean above, at, and below the Seafloor. Microbiol. Mol. Biol. Rev. 75(2):361–422.

4. Boyd PW, Claustre H, Levy M, Siegel DA, & Weber T (2019) Multi-faceted particle pumps drive carbon sequestration in the ocean. Nature 568(7752):327–335.

5. Jiao N, et al. (2010) Microbial production of recalcitrant dissolved organic matter: long-term carbon storage in the global ocean. Nature Reviews Microbiology 8:593.

6. Jorgensen BB & Boetius A (2007) Feast and famine--microbial life in the deep-sea bed. Nat Rev Microbiol 5(10):770–781.

7. Bergauer K, et al. (2018) Organic matter processing by microbial communities throughout the Atlantic water column as revealed by metaproteomics. Proceedings of the National Academy of Sciences of the United States of America 115(3):E400–E408.

8. Fontanez KM, Eppley JM, Samo TJ, Karl DM, & DeLong EF (2015) Microbial community structure and function on sinking particles in the North Pacific Subtropical Gyre. Front Microbiol 6:469.

9. Rieck A, Herlemann DPR, Jürgens K, & Grossart H-P (2015) Particle-Associated Differ from Free-Living Bacteria in Surface Waters of the Baltic Sea. Frontiers in Microbiology 6.

10. López-Pérez M, Kimes NE, Haro-Moreno JM, & Rodriguez-Valera F (2016) Not All Particles Are Equal: The Selective Enrichment of Particle-Associated Bacteria from the Mediterranean Sea. Frontiers in Microbiology 7.

11. Salazar G, et al. (2016) Global diversity and biogeography of deep-sea pelagic prokaryotes. The ISME journal 10(3):596–608.

12. Shah Walter SR, et al. (2018) Microbial decomposition of marine dissolved organic matter in cool oceanic crust. Nature Geoscience 11(5):334–339.

13. Boeuf D, et al. (2019) Biological composition and microbial dynamics of sinking particulate organic matter at abyssal depths in the oligotrophic open ocean. Proceedings of the National Academy of Sciences:201903080.

14. Zhao ZH, Baltar F, & Herndl GJ (2020) Linking extracellular enzymes to phylogeny indicates a predominantly particle-associated lifestyle of deep-sea prokaryotes. Science Advances 6(16):10.

15. Soltwedel T, Guilini K, Sauter E, Schewe I, & Hasemann C (2017) Local effects of large food-falls on nematode diversity at an arctic deep-sea site: Results from an in situ experiment at the deep-sea observatory HAUSGARTEN. J. Exp. Mar. Biol. Ecol.

16. Goffredi SK & Orphan VJ (2010) Bacterial community shifts in taxa and diversity in response to localized organic loading in the deep sea. Environ. Microbiol. 12(2):344–363.

17. Harbour RP, et al. (2021) Biodiversity, community structure and ecosystem function on kelp and wood falls in the Norwegian deep sea. Mar. Ecol. Prog. Ser. 657:73–91.

18. Avcı B, Krüger K, Fuchs BM, Teeling H, & Amann RI (2020) Polysaccharide niche partitioning of distinct *Polaribacter* clades during North Sea spring algal blooms. The ISME journal 14(6):1369–1383.

19. Kalenitchenko D, et al. (2016) Ecological succession leads to chemosynthesis in mats colonizing wood in sea water. The ISME journal 10(9):2246–2258.

20. Grabowski E, Letelier RM, Laws EA, & Karl DM (2019) Coupling carbon and energy fluxes in the North Pacific Subtropical Gyre. Nat Commun 10(1):1895.

21. Arnosti C (2011) Microbial Extracellular Enzymes and the Marine Carbon Cycle. Annual Review of Marine Science 3(1):401–425.

22. Liu R, et al. (2018) Depth-Resolved Distribution of Particle-Attached and Free-Living Bacterial Communities in the Water Column of the New Britain Trench. Front Microbiol 9.

23. Dai M, Yin Z, Meng F, Liu Q, & Cai W-J (2012) Spatial distribution of riverine DOC inputs to the ocean: an updated global synthesis. Current Opinion in Environmental Sustainability 4(2):170–178.

24. Schlünz B & Schneider RR (2000) Transport of terrestrial organic carbon to the oceans by rivers: re-estimating flux- and burial rates. International Journal of Earth Sciences 88(4):599–606.

25. Bianchi TS (2011) The role of terrestrially derived organic carbon in the coastal ocean: A changing paradigm and the priming effect. Proceedings of the National Academy of Sciences 108(49):19473–19481.

26. West AJ, et al. (2011) Mobilization and transport of coarse woody debris to the oceans triggered by an extreme tropical storm. Limnol. Oceanogr. 56(1):77–85.

27. Wohl E & Iskin Emily P (2021) Damming the wood falls. Science Advances 7(50):eabj0988.

28. Lee H, et al. (2019) Sustained wood burial in the Bengal Fan over the last 19 My. Proceedings of the National Academy of Sciences 116(45):22518–22525.

29. Pop Ristova P, Bienhold C, Wenzhöfer F, Rossel PE, & Boetius A (2017) Temporal and Spatial Variations of Bacterial and Faunal Communities Associated with Deep-Sea Wood Falls. PLOS ONE 12(1):e0169906.

30. Goffredi SK, Wilpiszeski R, Lee R, & Orphan VJ (2008) Temporal evolution of methane cycling and phylogenetic diversity of archaea in sediments from a deep-sea whale-fall in Monterey Canyon, California. The ISME journal 2:204.

31. Treude T, et al. (2009) Biogeochemistry of a deep-sea whale fall: sulfate reduction, sulfide efflux and methanogenesis. Mar. Ecol. Prog. Ser. 382:1–21.

32. Baco AR & Smith CR (2003) High species richness in deep-sea chemoautotrophic whale skeleton communities. Mar. Ecol. Prog. Ser. 260:109–114.

33. Li J, et al. (2022) Unique deep-sea bacterial assemblages thriving on different organic matters delivered via in-situ incubators. Preprint.

34. Watanabe M, Kojima H, & Fukui M (2020) *Labilibaculum antarcticum* sp. nov., a novel facultative anaerobic, psychrotorelant bacterium isolated from marine sediment of Antarctica. Antonie Van Leeuwenhoek 113(3):349–355.

35. Vandieken V, Marshall IPG, Niemann H, Engelen B, & Cypionka H (2018) *Labilibaculum manganireducens* gen. nov., sp. nov. and *Labilibaculum filiforme* sp. nov., Novel *Bacteroidetes* Isolated from Subsurface Sediments of the Baltic Sea. Frontiers in Microbiology 8(2614).

36. Wu W-J, Zhao J-X, Chen G-J, & Du Z-J (2016) Description of *Ancylomarina subtilis* gen. nov., sp. nov., isolated from coastal sediment, proposal of Marinilabiliales ord. nov. and transfer of *Marinilabiliaceae*, *Prolixibacteraceae* and *Marinifilaceae* to the order Marinilabiliales. Int. J. Syst. Evol. Microbiol. 66(10):4243–4249.

37. Xu Z-X, Mu X, Zhang H-X, Chen G-J, & Du Z-J (2016) Marinifilum albidiflavum sp. nov., isolated from coastal sediment. Int. J. Syst. Evol. Microbiol. 66(11):4589–4593.

38. Kalenitchenko D, et al. (2017) Bacteria alone establish the chemical basis of the wood-fall chemosynthetic ecosystem in the deep-sea. The ISME journal.

39. Kappelmann L, et al. (2019) Polysaccharide utilization loci of North Sea *Flavobacteriia* as basis for using SusC/D-protein expression for predicting major phytoplankton glycans. The ISME journal 13(1):76–91.

40. Krüger K, et al. (2019) In marine Bacteroidetes the bulk of glycan degradation during algae blooms is mediated by few clades using a restricted set of genes. The ISME journal 13(11):2800–2816.

41. Foley MH, Martens EC, & Koropatkin NM (2018) SusE facilitates starch uptake independent of starch binding in B. thetaiotaomicron. Mol. Microbiol. 108(5):551–566.

42. Najmudin S, et al. (2010) Putting an N-terminal end to the Clostridium thermocellum xylanase Xyn10B story: Crystal structure of the CBM22-1– GH10 modules complexed with xylohexaose. Journal of Structural Biology 172(3):353–362.

43. Rashid GMM, et al. (2015) Identification of Manganese Superoxide Dismutase from *Sphingobacterium* sp. T2 as a Novel Bacterial Enzyme for Lignin Oxidation. ACS Chemical Biology 10(10):2286–2294.

44. Liu X, et al. (2017) Crystal structure and biochemical features of dye-decolorizing peroxidase YfeX from *Escherichia coli* O157 Asp143 and Arg232 play divergent roles toward different substrates. Biochem. Biophys. Res. Commun. 484(1):40–44.

45. Rashid GMM, et al. (2018) *Sphingobacterium* sp. T2 Manganese Superoxide Dismutase Catalyzes the Oxidative Demethylation of Polymeric Lignin via Generation of Hydroxyl Radical. ACS Chem Biol 13(10):2920–2929.

46. Rodríguez-Beltrán J, Rodríguez-Rojas A, Guelfo JR, Couce A, & Blázquez J (2012) The Escherichia coli SOS gene dinF protects against oxidative stress and bile salts. PLoS One 7(4):e34791.

47. Ahmad M, et al. (2011) Identification of DypB from *Rhodococcus jostii* RHA1 as a Lignin Peroxidase. Biochemistry 50(23):5096–5107.

48. Voight JR (2015) Xylotrophic bivalves: aspects of their biology and the impacts of humans. Journal of Molluscan Studies 81(2):175–186.

49. Hedges JI, et al. (2001) Evidence for non-selective preservation of organic matter in sinking marine particles. Nature 409(6822):801–804.

50. Veith PD, Glew MD, Gorasia DG, & Reynolds EC (2017) Type IX secretion: the generation of bacterial cell surface coatings involved in virulence, gliding motility and the degradation of complex biopolymers. 106(1):35–53.

51. Hu Y & Ribbe MW (2016) Biosynthesis of the Metalloclusters of Nitrogenases. Annu. Rev. Biochem. 85(1):455–483.

52. Sickerman NS, Hu Y, & Ribbe MW (2019) Nitrogenases. Metalloproteins: Methods and Protocols, ed Hu Y (Springer New York, New York, NY), pp 3–24.

53. Inoue J-I, et al. (2015) Distribution and Evolution of Nitrogen Fixation Genes in the Phylum Bacteroidetes. Microbes and environments / JSME 30:44–50.

54. Lilburn TG, et al. (2001) Nitrogen fixation by symbiotic and free-living spirochetes. Science 292(5526):2495–2498.

55. Suzuki Y, et al. (2009) Molecular investigations of the stalked barnacle Vulcanolepas osheai and the epibiotic bacteria from the Brothers Caldera, Kermadec Arc, New Zealand pp 727–733.

56. Bik EM, et al. (2016) Marine mammals harbor unique microbiotas shaped by and yet distinct from the sea. Nature communications 7:10516.

57. Sunagawa S, et al. (2009) Bacterial diversity and White Plague Disease-associated community changes in the Caribbean coral Montastraea faveolata. The ISME journal 3:512.

58. Kapili BJ, Barnett SE, Buckley DH, & Dekas AE (2020) Evidence for phylogenetically and catabolically diverse active diazotrophs in deep-sea sediment. The ISME journal 14(4):971–983.

59. Lloyd JR (2003) Microbial reduction of metals and radionuclides. FEMS Microbiol. Rev. 27(2-3):411–425.

60. Shi L, et al. (2016) Extracellular electron transfer mechanisms between microorganisms and minerals. Nature Reviews Microbiology 14(10):651–662.

61. Greening C, et al. (2016) Genomic and metagenomic surveys of hydrogenase distribution indicate H-2 is a widely utilised energy source for microbial growth and survival. Isme Journal 10(3):761–777.

62. Stramma L, Johnson GC, Sprintall J, & Mohrholz V (2008) Expanding Oxygen-Minimum Zones in the Tropical Oceans. Science 320(5876):655–658.

63. Matear RJ & Hirst AC (2003) Long-term changes in dissolved oxygen concentrations in the ocean caused by protracted global warming. Global Biogeochemical Cycles 17(4).

64. Dong X, et al. (2020) Thermogenic hydrocarbon biodegradation by diverse depth-stratified microbial populations at a Scotian Basin cold seep. Nature communications 11(1):5825.

65. Dong X, et al. (2019) Metabolic potential of uncultured bacteria and archaea associated with petroleum seepage in deep-sea sediments. Nature communications 10(1):1816.

66. Chen S, Zhou Y, Chen Y, & Gu J (2018) fastp: an ultra-fast all-in-one FASTQ preprocessor. Bioinformatics 34(17):i884–i890.

67. Langmea d B & Salzberg SL (2012) Fast gapped-read alignment with Bowtie 2. Nat. Methods 9(4):357–359.

68. Nurk S, Meleshko D, Korobeynikov A, & Pevzner PA (2017) metaSPAdes: a new versatile metagenomic assembler. Genome Res. 27(5):824–834.

69. Uritskiy GV, DiRuggiero J, & Taylor J (2018) MetaWRAP—a flexible pipeline for genome-resolved metagenomic data analysis. Microbiome 6(1):158.

70. Parks DH, Imelfort M, Skennerton CT, Hugenholtz P, & Tyson GW (2015) CheckM: assessing the quality of microbial genomes recovered from isolates, single cells, and metagenomes. Genome Res.

71. Chaumeil P-A, Mussig AJ, Hugenholtz P, & Parks DH (2019) GTDB-Tk: a toolkit to classify genomes with the Genome Taxonomy Database. Bioinformatics 36(6):1925–1927.

72. Jain C, Rodriguez-R LM, Phillippy AM, Konstantinidis KT, & Aluru S (2018) High throughput ANI analysis of 90K prokaryotic genomes reveals clear species boundaries. Nature communications 9(1):5114.

73. Hyatt D, et al. (2010) Prodigal: prokaryotic gene recognition and translation initiation site identification. BMC Bioinformatics 11(1):119.

74. Moriya Y, Itoh M, Okuda S, Yoshizawa AC, & Kanehisa M (2007) KAAS: an automatic genome annotation and pathway reconstruction server. Nucleic Acids Res. 35(Web Server issue):W182–185.

75. Huerta-Cepas J, et al. (2017) Fast Genome-Wide Functional Annotation through Orthology Assignment by eggNOG-Mapper. Mol. Biol. Evol. 34(8):2115–2122.

76. El-Gebali S, et al. (2018) The Pfam protein families database in 2019. Nucleic Acids Res. 47(D1):D427–D432.

77. Rawlings ND, et al. (2017) The MEROPS database of proteolytic enzymes, their substrates and inhibitors in 2017 and a comparison with peptidases in the PANTHER database. Nucleic Acids Res. 46(D1):D624–D632.

78. Zhang H, et al. (2018) dbCAN2: a meta server for automated carbohydrate-active enzyme annotation. Nucleic Acids Res. 46(W1):W95–W101.

79. Buchfink B, Xie C, & Huson DH (2014) Fast and sensitive protein alignment using DIAMOND. Nat. Methods 12:59.

80. Kelley LA, Mezulis S, Yates CM, Wass MN, & Sternberg MJE (2015) The Phyre2 web portal for protein modeling, prediction and analysis. Nature Protocols 10(6):845–858.

81. Liao Y, Smyth GK, & Shi W (2013) featureCounts: an efficient general purpose program for assigning sequence reads to genomic features. Bioinformatics 30(7):923–930.

82. Wagner GP, Kin K, & Lynch VJ (2012) Measurement of mRNA abundance using RNA-seq data: RPKM measure is inconsistent among samples. Theory in biosciences = Theorie in den Biowissenschaften 131(4):281–285.

83. Kumar S, Stecher G, & Tamura K (2016) MEGA7: Molecular Evolutionary Genetics Analysis Version 7.0 for Bigger Datasets. Mol. Biol. Evol. 33(7):1870–1874.

84. Na S-I, et al. (2018) UBCG: Up-to-date bacterial core gene set and pipeline for phylogenomic tree reconstruction. Journal of Microbiology 56(4):280–285.

